# Using EEG to characterise drowsiness during short duration exposure to elevated indoor Carbon Dioxide concentrations

**DOI:** 10.1101/483750

**Authors:** Stephen Snow, Amy Boyson, Marco Felipe-King, Obaid Malik, Louise Coutts, Catherine J Noakes, Hannah Gough, Janet Barlow, m.c. schraefel

## Abstract

Drowsiness which can affect work performance, is often elicited through self-reporting. This paper demonstrates the potential to use EEG to objectively quantify changes to drowsiness due to poor indoor air quality. Continuous EEG data was recorded from 23 treatment group participants subject to artificially raised indoor CO_2_ concentrations (average 2,700 ± 300 ppm) for approximately 10 minutes and 13 control group participants subject to the same protocol without additional CO_2_ (average 830 ± 70 ppm). EEG data were analysed for markers of drowsiness according neurophysiological methods at three stages of the experiment, Baseline, High CO_2_ and Post-Ventilation. Treatment group participants’ EEG data yielded a closer approximation to drowsiness than that of control group participants during the High CO_2_ condition, despite no significant group differences in self-reported sleepiness. Future work is required to determine the persistence of these changes to EEG over longer exposures and to better isolate the specific effect of CO_2_ on drowsiness compared to other environmental or physiological factors.

**Practical implications:**

- This study introduces EEG as a potential objective indicator of the effect of indoor environmental conditions upon drowsiness
- Participants exposed to 2,700 ppm for 10 minutes showed greater evidence of a progression towards drowsiness (as measured by EEG) than that of participants who received the same protocol without additional CO_2_ (mean 830 ± 70 ppm), despite similar ratings of subjective sleepiness.
- Subjective and objectively measured indications of drowsiness were reduced following ventilation of the room. Future work could explore the potential of regular ventilation episodes in knowledge work spaces to retain alertness.

## Introduction

Being a product of human respiration, carbon dioxide (CO_2_) increases in indoor spaces when ventilation of the space is insufficient to replace stale air [1,2]. CO_2_ is thus a useful indicator of ventilation and, by extension air quality indoors, in occupied spaces [3,4]. A large body of literature exists relating poor ventilation to mild health symptoms [2,5-7] and lowered cognitive performance [4,8-10]. Office-realistic levels of CO_2_ are reported to be typically < 3,000 ppm, but whether CO_2_ itself negatively impacts cognitive performance, or whether other pollutants such as volatile organic compounds (VOCs), including human bio-effluents, are responsible, is still unclear [11,12]. Human performance effects have been recorded in studies both where CO_2_ is accompanied by human bio-effluents (e.g. the CO_2_ concentration is a product of poor ventilation in occupied spaces) [4,13,14] and where pure CO_2_ gas is added to a room to achieve steady-state concentrations [12,13,15–18].

At a room concentration of 3,000 ppm, human bio-effluents are found to cause an increase in respired -end-tidal- CO_2_ (ETCO_2_), increased blood pressure, and seemingly increased stress/arousal, as well as reduced cognitive performance [19]. Zhang et al. proposes CO_2_ with bio-effluents affects cognitive performance through either (1) stress/arousal or (2) physiological factors such as an increase in ETCO_2_ and reduced nasal peak flow, triggering symptoms such as subjective (self-reported) sleepiness, tiredness and headache [19]. One study found that four hours of exposure to non-ventilated conditions, with average CO_2_ concentrations above 2,700 ppm, resulted in significantly increased blood-CO_2_, heart rate variability, and increased peripheral blood flow, as well as increased prevalence of health symptoms and self-reported sleepiness [14]. The study reports that the high CO_2_ concentration itself (separate to bio-effluents) may be a parameter affecting physiology which can lower functional ability and increase (self-reported) sleepiness [14]. Given findings that 1,400 ppm [15] and 2,500 ppm [16] of CO_2_ achieved by introducing pure gas into a room correlates to lower decision making capability, cumulatively, there is some evidence that CO_2_ itself, independent of other indoor pollutants, may play a role in detrimentally affecting aspects of work performance [15–17].

Drowsiness and fatigue are recognised as important parameters affecting office work and productivity [14,20]. In this study we focus on drowsiness, (i.e. lethargy or wish to sleep) [21–23], rather than mental fatigue (i.e. exhaustion or lack of motivation for task(s) due to extended work effort) [24]. Sub-optimal air quality (i.e. poor ventilation/high CO_2_) is correlated to increased self-reported sleepiness and fatigue [14]. Yet factors such as sleepiness, drowsiness and fatigue, when reported in studies assessing the effect of indoor conditions on humans, are often elicited subjectively through questionnaires only [10,14,18,25]. One study uses voice analysis to as a means of objectively measuring fatigue [20], but this method has not been widely adopted. The lack of objective measurement of drowsiness or fatigue may be problematic, given that self-reporting is identified as a less reliable measurement than objective measurement [26,27]. On the other hand, fields such as Neurophysiology, have a long history of objectively measuring sleep, and wakeful sleepiness/drowsiness using electroencephalogram (EEG). EEG records electrical activity in the brain using electrodes fitted to a cap, or placed on the scalp directly [28]. EEG data can be analysed to: (a) detect specific events (event-related potential) or (b) time-averaged power in different frequency bands [28]. A dominance of low frequency power is typically associated with lower neurological arousal (delta, theta) [22].

The impact of office-realistic concentrations of CO_2_ upon objectively measured drowsiness is a knowledge gap in the literature. Temperature effects on drowsiness using EEG find lower temperatures are correlated to reduced drowsiness [29]. EEG research to date concentrates on neurological effects of much higher concentrations of CO_2_ than is likely to occur in indoor spaces, e.g. 5% CO_2_/air mixture (50,000 ppm) [30–32], or 10% (100,000 ppm) [33] and the resultant hypercapnia (elevated blood CO_2_) [30,31,33]. In these studies EEG results are assessed according to arousal state (i.e. overall changes to low-frequency parameters), but not drowsiness specifically. Xu et al.[31] found inhalation of a 5% CO_2_/air mixture (50,000 ppm) caused transition to a lower (brain) arousal state, characterised by a relative increase in delta power and corresponding decrease in alpha power. Bloch-Salisbury [33] subjected participants to 10% CO_2_ (10,000 ppm) through direct inhalation, finding a significant decrease in both overall power and a movement of the centroid frequency (i.e. the centrepoint of the mass of frequencies observed) toward lower frequencies.

In summary, (1) findings are mixed as to whether CO_2_ is a pollutant affecting cognitive performance in its own right, with some studies finding evidence that CO_2_ affects cognitive performance [15–17], while others find no evidence of this relationship [10,13,18]. (2) Poor indoor air quality is correlated to increased subjective drowsiness [14], yet drowsiness is typically elicited through self-reporting [10,14,18,25], which is less reliable than objective measurement [26,27]. (3) the field of neurophysiology offer methods of objectively measuring drowsiness (a precursor to sleepiness) using EEG [23,34], yet these methods have not yet been applied to office-realistic CO_2_ concentrations. (4) Literature on the effect of CO_2_ on resting EEG is presently limited to the human effects of much higher levels of CO_2_ [31,33] than could realistically be achieved indoors through human respiration. Comparable studies of office- realistic concentrations of CO_2_ are not yet available, providing impetus for this present paper.

This paper details the novel application of using electroencephalogram (EEG) as a means of objectively measuring the effect of CO_2_ on drowsiness at office-realistic concentrations. Resting EEG and other physiological and subjective parameters were recorded from participants exposed to 2,700 ± 300 ppm of CO_2_ in an office for 10 minutes, as a means of determining the physiological changes of a short-duration exposure to elevated CO_2_ concentration and testing for EEG data indicative of a progression towards drowsiness. A key aim of the paper is to explore the effect of CO_2_ on drowsiness, given that drowsiness is a determinant of human work performance [20,35] and compare results to both cognitive science literature on the cognitive performance effects of office-realistic concentrations of CO_2_ [4,8,10,12,15,16] and neurophysiology literature on the neurological effects of much higher concentrations of CO_2_ [30–33].

## Materials and methods

### Rationale for study design

Our chosen target for CO_2_ concentration (2,700ppm) reflects a high, but realistic level achieved in occupied spaces when windows and doors are closed [2,14]. In a meta-review of classroom ventilation, Fisk [2] found six studies of 20 or more classrooms recorded average or median CO_2_ concentrations between 2,000 and 3,000 ppm. The target concentration is chosen to be comparable with other studies assessing the human performance effects of indoor CO_2_ concentration, e.g. 2,260 ppm [4] 2,500 ppm [16] or 3,000 ppm [10,17,19]. The duration of exposure to elevated CO_2_ concentration in our study is shorter compared to others [4,14,16], and relates to our aim to record and analyse EEG continuously throughout the experiment to provide a novel focus on immediate-term physiological effects of CO_2_. Continuous EEG recording is less practicable over extended study durations due to the need for participants to remain still during EEG recordings to ensure clean data [28]. The need to remain still over extended durations, when combined with a lack of stimulation may produce a tendency to fidget, which may in turn affect measured EEG parameters, or potentially cause boredom/ drowsiness itself, which could confound determination of drowsiness as caused through changing indoor environment parameters.

### Participants

A total of 47 subjects were recruited and participated in the study between October 2016 and February 2017. Usable EEG data was available from 36 of the 47 participants, reflective of the sensitivity of EEG to movement artefacts and the researchers’ wish for data reliability. The study protocol and conditions of participation were approved by the University of Southampton Ethical Research Governance Office (ERGO# 30443). Sampling was achieved by advertising the study on billboards throughout the University, a local supermarket and a departmental mailing list. Convenience sampling was used for contacts of the research team who were unaware of the study protocol. The final sample was comprised mostly of students and staff from the University. Written consent was gathered from each participant prior to their participation in the study. Exclusion criteria for the study were adapted from those used by Garner et al. [36], a study where participants were subjected to 7.5% CO_2_ (75,000 ppm) level of CO_2_. Exclusion criteria included current or historic drug/alcohol abuse or panic attacks, current treatment for migraine headaches, pregnant, current neurological conditions (e.g. epilepsy), and recent severe illness. Participants were compensated £10 in vouchers for an online retailer for their participation.

Participants were split into two groups. Of the participants with usable EEG data, this involved:23 participants in the “treatment group” (TG) who received artificially raised CO_2_ concentrations and 13 participants in the “control group” (CG) for whom CO_2_ concentrations were not artificially raised (Table 1). The variance in the size of the groups is due to which of the participants had sufficiently clean EEG for inclusion and the difficulty in recruiting a larger sample.

**Table 1:**
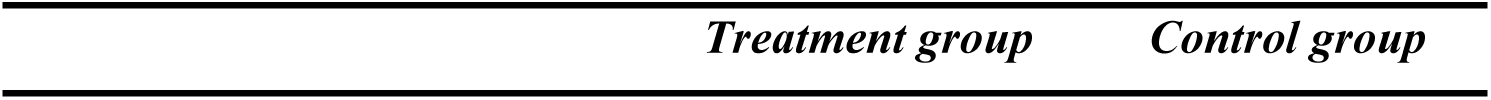
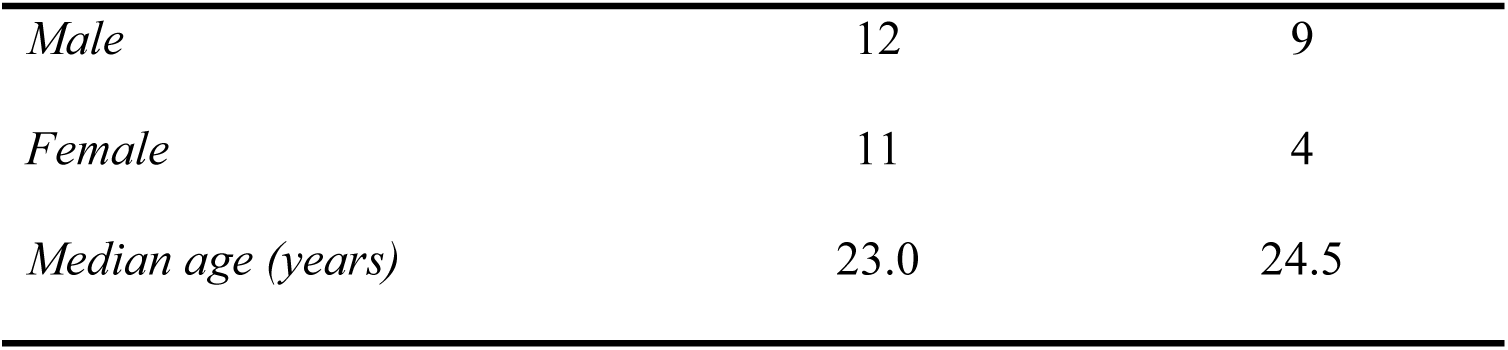
Participant attributes

| *Treatment group* | *Control group* |
| --- | --- |

|  |  |  |
| --- | --- | --- |
| *Male* | 12 | 9 |
| *Female* | 11 | 4 |
| *Median age (years)* | 23.0 | 24.5 |

Statistical power analysis was calculated a-priori using G*Power software [37]. Effect size was estimated at 0.4 based on similar experiments [12], number of groups = 2 (treatment, control), number of measurements = 3 (Baseline, High-CO_2_, Post ventilation- defined below), significance level 0.05. This gave a between factors recommendation for 58 participants, and recommendations for both within-factors and within-between factors of 18 participants. In this paper we concentrate on within-factors analysis.

### Study room

A motivation for the study was to replicate office-realistic scenarios. All experiments took place in a small, carpeted, naturally ventilated office of dimensions 4,000 mm by 3,400 mm (floor area) by 3,050 mm (high) (Figure 1). The office was on the fourth floor, on the northern end of a large building in the south of England. The office had two high windows on the north and west corner of the room. Only the western window could be opened, and is visible behind the participant in Figure 1. The CO_2_ cylinder was positioned directly in front of the openable window. The door of the room led to a larger reception office which was occupied by one staff member during some but not all of the experiments. The numbered arrows in Figure 1 point to the location of the CO_2_ loggers.

**Figure 1:**
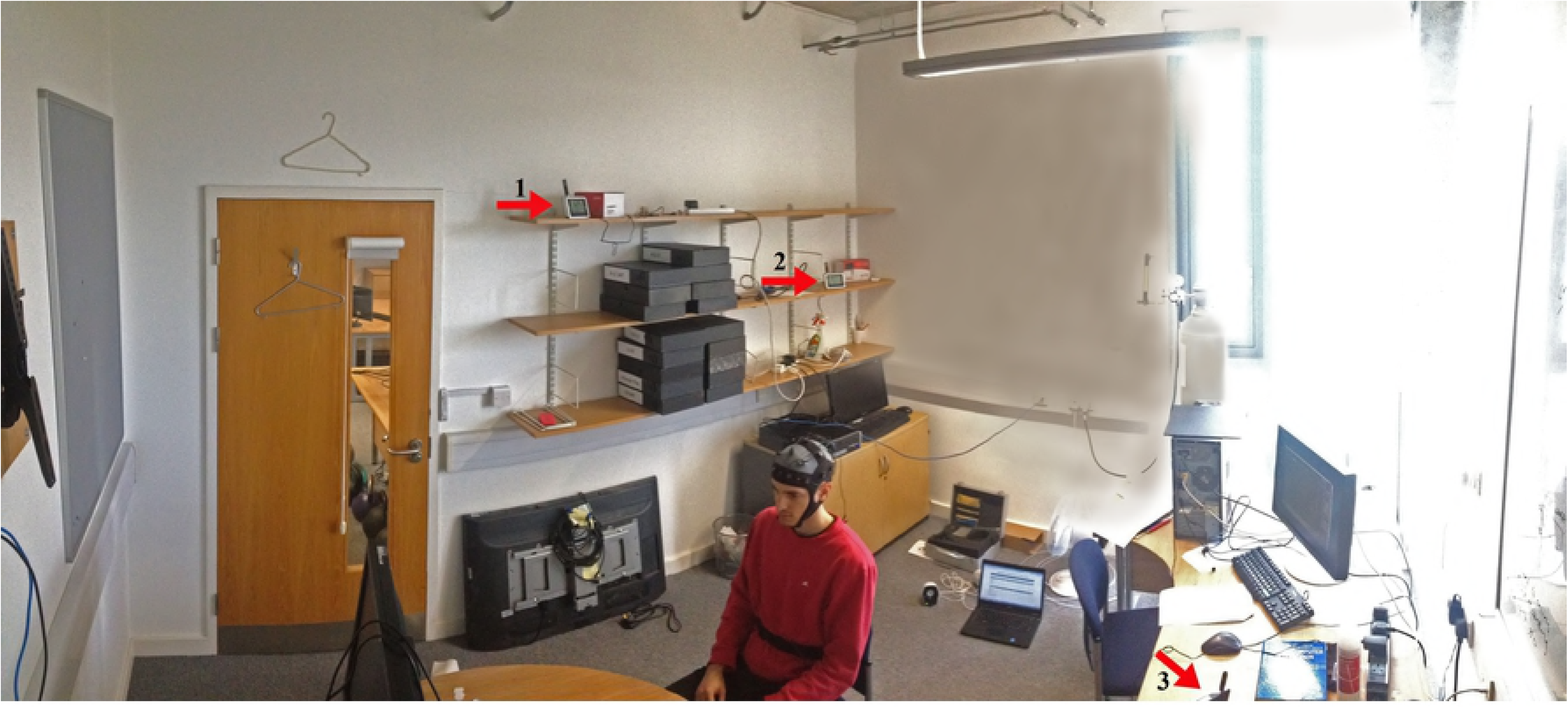
Study room showing participant with EEG cap, location of loggers, window and CO_2_ cylinder.

The infiltration rate of the study room with the windows closed was calculated according to Laussmann et al. [38] using a tracer-gas decay method overnight, with the researcher ensuring the mixing of CO_2_ in the room by observing the range of the readings from the three CO_2_ monitors and ensuring all were within instrument error before leaving the room overnight. This method gave an infiltration rate of 0.078 ± 0.002 (R^2^ = 0.91) air changes per hour, consistent with the rubber-sealed windows and minimal air gaps around the door. The value is approximate, given air exchange rates can differ over time due to differences in temperature, wind direction and wind speed [39].

Carbon dioxide was introduced using a cylinder of ultrapure CO_2_ (greater than 99.99% purity) located in the corner of the room with the outlet attached to pedestal fan to achieve mixing. The fan was pointed away from the participant and in operation only for the duration of Condition 3 (see Table 2), when CO_2_ was being released, in order to minimise any influence of air movement on perception or produce possible thermal comfort effects during subsequent conditions. The target CO_2_ concentration once mixed was 2,700 ppm (mean: 2,700 ± 300 ppm for the duration of Condition 5). Participants were instructed to sit at the table in the middle of the room while the researcher operated the computer and the gas cylinder behind the participant. In this way participants were aware the air quality was going to be changed somehow during the experiment, but were not aware how.

**Table 2:**
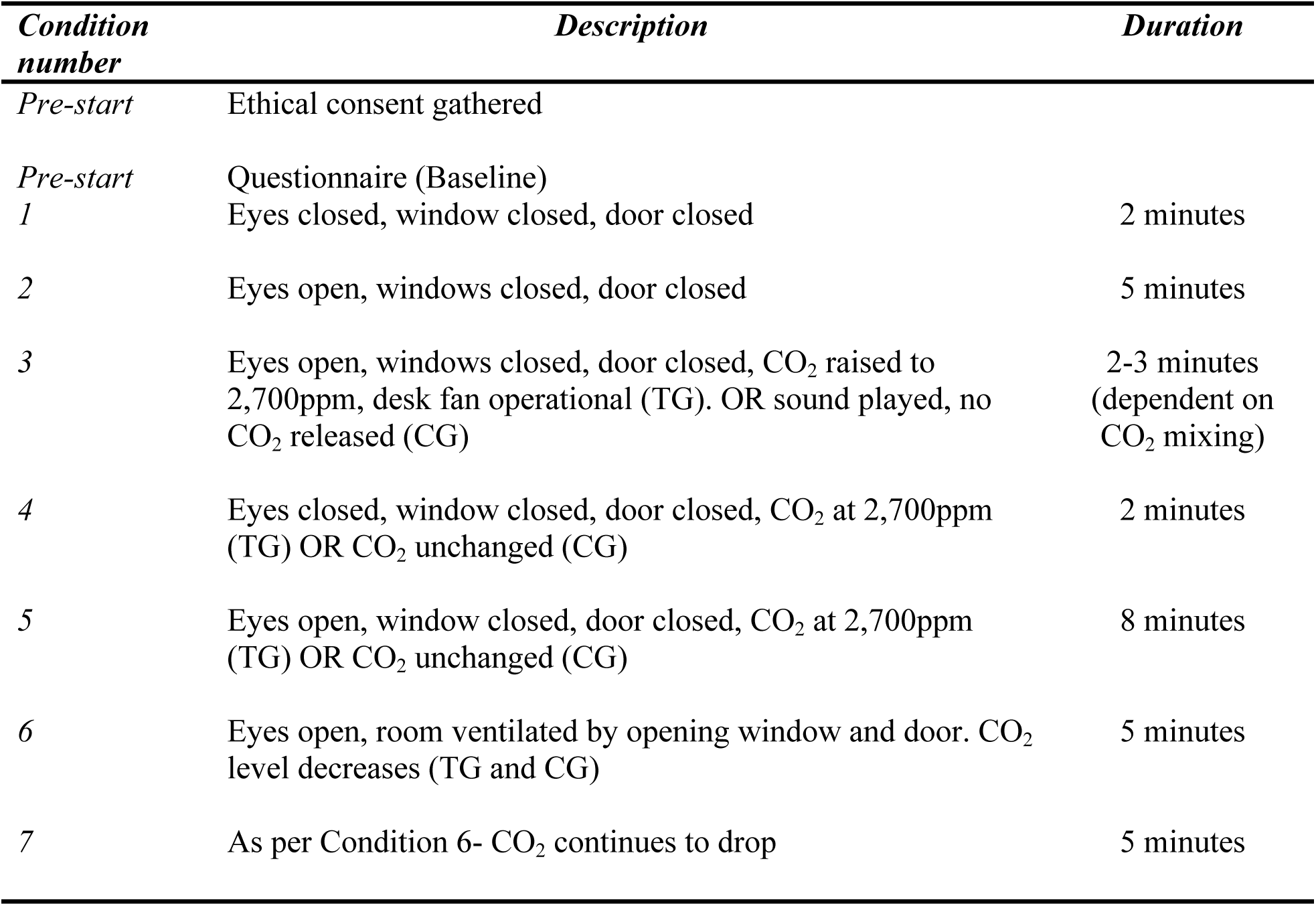
Experimental protocol

| **Condition number** | **Description** | **Duration** |
| --- | --- | --- |
| *Pre-start* | Ethical consent gathered |  |
| *Pre-start* | Questionnaire (Baseline) |  |
| 1 | Eyes closed, window closed, door closed | 2 minutes |
| 2 | Eyes open, windows closed, door closed | 5 minutes |
| 3 | Eyes open, windows closed, door closed, CO<sub>2</sub> raised to 2,700ppm, desk fan operational (TG). OR sound played, no CO<sub>2</sub> released (CG) | 2-3 minutes (dependent on CO<sub>2</sub> mixing) |
| 4 | Eyes closed, window closed, door closed, CO<sub>2</sub> at 2,700ppm (TG) OR CO<sub>2</sub> unchanged (CG) | 2 minutes |
| 5 | Eyes open, window closed, door closed, CO<sub>2</sub> at 2,700ppm (TG) OR CO<sub>2</sub> unchanged (CG) | 8 minutes |
| 6 | Eyes open, room ventilated by opening window and door. CO<sub>2</sub> level decreases (TG and CG) | 5 minutes |
| 7 | As per Condition 6- CO<sub>2</sub> continues to drop | 5 minutes |

### Experimental Procedure

The experimental protocol took place in the one study room (Figure 1). The study protocol is summarised below for TG participants (Table 2). CG participants experienced the same protocol to that of TG participants, except that the CO_2_ concentration of the room was not modified using the cylinder. Instead a pre-recorded and equalized sound was used in place of the CO_2_ gas being released throughout Condition 3 to mimic the sound of the gas release. When questioned, no CG participant identified the sound as audio playback and thus every participant assumed their air quality was being modified.

For comparative data analysis, three two-minute segments were selected for comparison, (1) Baseline – the first two minutes of Condition 2; before the environmental conditions were changed, (2) High-CO_2_ – The last two minutes of Condition 5; beginning when TG participants had been exposed to the higher CO_2_ concentration for 8 minutes; and (3) Post-Ventilation - last two minutes of Condition 7, beginning after 8 minutes of room ventilation. The location of these analysis segments within the context of the study protocol are shown in Figure 2.

**Figure 2:**
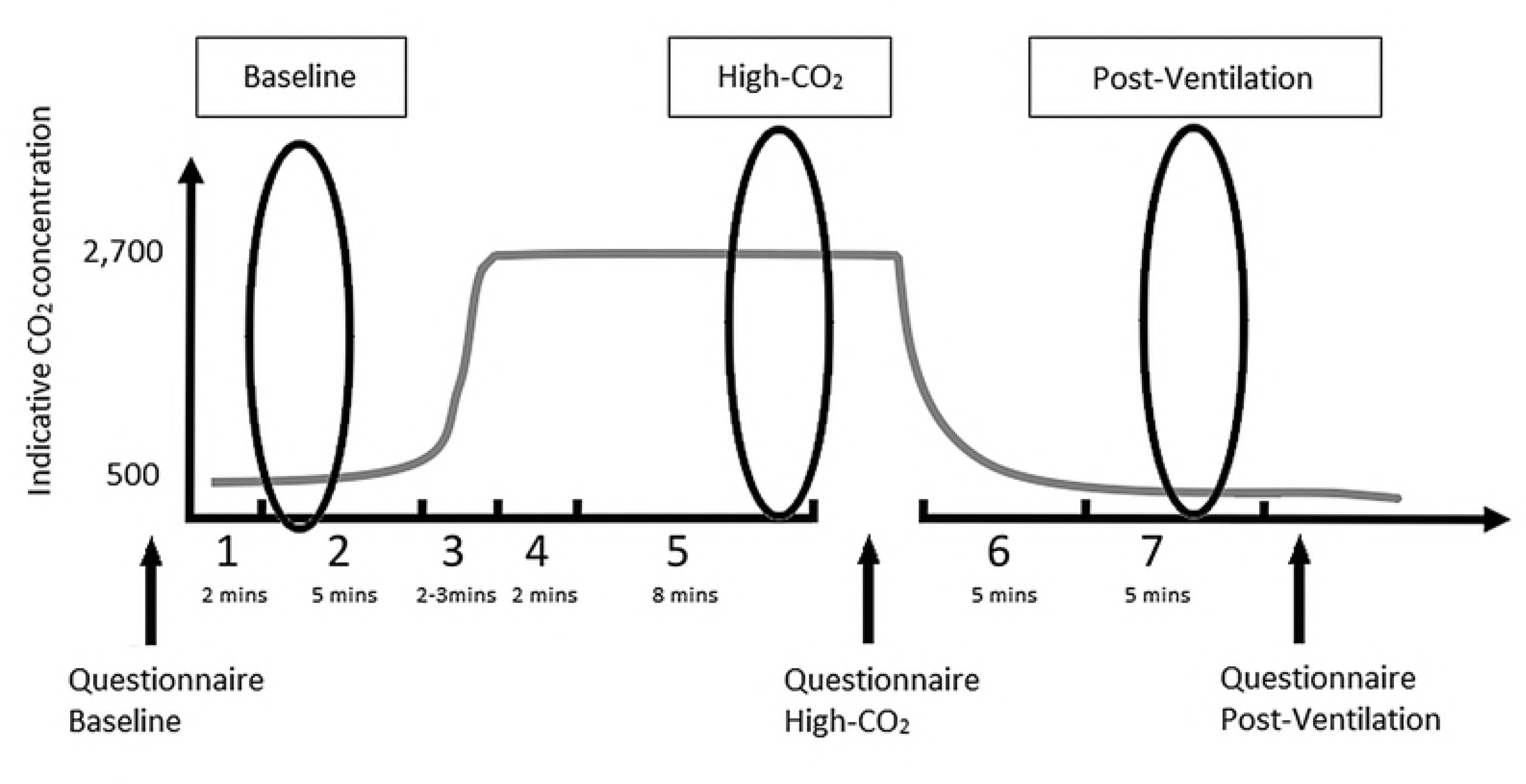
Study protocol with indicative CO_2_ level showing location of Baseline, High-CO_2_ and Post-Ventilation segments.

### Measurement

Three factory calibrated Rotronic CL11 (BSRIA, Bracknell, UK) environmental loggers measured temperature, humidity and CO_2_ concentration throughout each experiment. The loggers were positioned approximately equidistant around the room and are labelled 1, 2 and 3 in Figure 1. The loggers were positioned so as to avoid influence from direct respiration. The heights of the loggers from the floor were 720 mm (logger 1), 1,545 mm (logger 2) and 1,995 mm (logger 3). The distance from logger 2 to logger 3 was 2,100 mm and logger 1 was approximately 1,300 mm perpendicular to the participant’s heads (Figure 1). Instrument accuracies for the CL11 are ± 0.3 ° C (temperature), < 2.5% RH (humidity) and ± 30 ppm ± 5% of the measured value. The logging frequency of the CL11 monitors was set to 10 seconds throughout the experiments. The CL11’s display updates approximately once per second, enabling the researcher to monitor and control the release of CO_2_ in the room to a reasonable granularity. The length of Condition 3 (adding CO_2_) was varied according to the time taken to achieve mixing (Table 2), to enable confidence in the mixing of the room by the start of Condition 4.

EEG data was gathered from each participant using a Neuroelectrics ENOBIO 20 dry electrode wearable wireless EEG cap (19 channel, 10-20 placement, 500 Hz sampling rate). Two reference electrodes (DLR, CRL) were positioned on the participants’ mastoid muscle. EEG was gathered continuously throughout each of the experimental conditions (Table 2, Figure 2). In order to minimise movement artefacts in the EEG, participants were asked to sit quietly and remain still throughout the experiments except during the short break for the questionnaire following Condition 5 (refer Table 2, Figure 2).

Subjective responses were gathered in relation to experience of sick building symptoms (e.g. irritated eyes, sore throat, congested nose) [40], positive/negative affect (PANAS) [41], Stanford Sleepiness Scale [42] and thermal comfort (ASHRAE 7 point scale) [43] were gathered from participants at Baseline, High- CO_2_ and Post-Ventilation segments.

As a proof-of-concept, this paper focuses specifically on EEG results and the Stanford Sleepiness Scale.

### Analysis

#### Environmental measurements

Data from the Rotronic CL11 environmental monitors was downloaded and condition timings entered retrospectively for analysis. Due to the difference in logging frequency of the CL11s (10 sec) compared to the EEG measurements (500 Hz), the error on the readings versus that of the condition timings is expected to be approximately ± 20 seconds. This error was considered acceptable given the gradual changes in temperature/humidity and the mixing behaviour of the CO_2_ in the room.

#### EEG pre-processing

EEG data were filtered using a Butterworth filter; low pass at 45 Hz and high pass at 0.15 Hz. Artefact rejection was implemented in two stages. The first used the artefact rejection algorithm WPT-EMD [44,45], which uses a sample of minimum variance EEG taken from Condition 2. The second stage of artefact rejection involved an amplitude threshold cut-off of ±100 μV, and replacing outlying data with a 10-second moving median around the extreme value. Electrodes showing consistent noise or flat-lined output were deleted from the dataset. As mentioned, of the total 47 participants, 36 participants had sufficiently clean data throughout the experiment and sufficient representation of clean electrodes in each brain region (frontal, central, temporal, parietal, occipital) to warrant further analysis.

Bandpower was extracted from the pre-processed continuous EEG for delta (0.15-3 Hz), theta (4-7 Hz), alpha (8-13 Hz), beta (14-35 Hz), and gamma (> 35 Hz) frequency bands, over one second windows. Average bandpower was computed for frontal (F3, Fz, F4, FP1, FP2), central (C3, Cz, C4), parietal (P7, P3, Pz, P4, P8), temporal (T7, T8), and occipital (O1, O2) electrodes for each analysis segment (Baseline, High-CO_2_, Post-Ventilation). Gamma was excluded from further analysis owing to the focus of the study protocol on low frequency behaviour and because gamma represented < 1% of total power at each analysis segment. Post-hoc analysis found the lowest delta component (0.15-1.5 Hz) to be contaminated with eye movement artefacts and was subsequently rejected from analysis. Rather than excluding delta from analysis completely, and given eye movement artefacts typically occur at approximately 1 Hz [46], we instead report on high-delta (2-3 Hz) and exclude only low-delta (i.e. all frequencies < 2 Hz).

Mixed model ANOVAs were conducted with factors including electrode region, analysis segment, group, and frequency to investigate electrophysiological markers of drowsiness consistent with the literature (detailed below).

#### EEG- drowsiness characterisation

Our characterisation of drowsiness applied to the EEG results is grounded in relevant literature: A meta-review of the psychophysiology of automobile driver fatigue finds changes in delta and theta strongly linked to the transition towards fatigue [21]. Tired wakefulness among sleep deprived participants produces an EEG with enhanced power in the low frequency range 1-8 Hz (delta and theta) [22,47]. Providing a greater topographical specificity than previous studies, Gorgoni et al. finds sleep deprived participants exhibit an EEG involving global increases in delta and theta (i.e. registered in multiple areas of the brain) [23]. Thus in this study, drowsiness is characterised by post-hoc analysis of the cleaned EEG data according to an increase in delta and theta, particularly if these increases are found at multiple brain electrode regions.

## Results

All statistical analyses conducted and reported in this section relate to data from the three analysis segments of Baseline, High-CO_2_ and Post-Ventilation. Additionally, all analyses and data reported below relate to the 36 participants with usable EEG data.

### Indoor conditions by analysis segment

Table 3 below summarises the measured indoor environment parameters at each of the two-minute analysis segments: Baseline, High- CO_2_ and Post-Ventilation (Figure 2), for TG and CG participants:

**Table 3:** Indoor parameters by analysis segment, TG and CG participants

| *Treatment group participants* | *CO<sub>2</sub> (ppm)* | *Temp (°C)* | *RH<sup>1</sup> (%)* |
| --- | --- | --- | --- |
| *Baseline* | 670 ± 80 | 21.8 ± 2.3 | 44.1 ± 8.2 |
| *High-CO<sub>2</sub>* | 2750 ± 160 | 22.2 ± 2.5 | 44.6 ± 7.9 |
| *Post-Ventilation* | 850 ± 210 | 21.5 ± 2.4 | 43.9 ± 7.5 |
| *Control group participants* | *CO<sub>2</sub> (ppm)* | *Temp (°C)* | *RH<sup>1</sup> (%)* |
| *Baseline* | 660 ± 40 | 23.6 ± 1.8 | 37.7 ± 7.6 |
| *High-CO<sub>2</sub>* | 860 ± 50 | 24.3 ± 2.0 | 37.8 ± 7.3 |
| *Post-Ventilation* | 680 ± 80 | 23.4 ± 1.8 | 37.8 ± 7.8 |

The mean CO_2_ values for the two minute segments of Baseline and High-CO_2_ correspond closely to the mean CO_2_ values for TG and CG participants for entire five minute duration of Condition 2 (650 ± 80 ppm TG, 640 ± 50 ppm CG) and eight minute duration of Condition 5 (2,700 ± 300 ppm in TG, 830 ± 70 in CG). With reference to Table 3, TG participants were exposed on average to an additional 1,898 ppm of pure CO_2_ to that generated by human respiration alone.

To control for possible temperature effects, all participants were able to adjust clothing as they wished prior to the experiment to ensure comfort. A 3 (analysis segment) by 2 (group) mixed model ANOVA was run to assess temperature fluctuations. Results show that CG participants were tested at a significantly higher temperature than TG participants (see Table 3 and Section 0; *F* (1, 34) = 6.30, *p* = .02,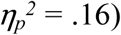 This was due to the majority of CG participants being tested following the activation of the building’s heating systems. Results also showed that temperature varied significantly between each of the analysis segments irrespective of group (*F* (1.46, 49.55) = 50.75, *p* < .001, 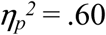Sidak post-hoc *p’s* < .02). Temperature was higher on average for both groups at High CO_2_ relative to the other conditions, due to the doors and windows remaining closed; additionally, Post-Ventilation was colder than both High CO_2_ and Baseline for both groups due to the windows being open throughout the condition and the cooler outside air due to the season. However, the difference in temperature between analysis segments (i.e. Baseline vs High CO_2_ vs Post-Vent) did not greatly exceed instrument accuracy (0.3 °C).

The period of ventilation (including the Post-Ventilation analysis segment) was uncontrolled. During this period, CO_2_ concentration (Table 3), as well as air change rate, indoor air velocity and external noise was variable between participants, depending on external factors such as wind direction, wind speed and traffic. We did not attempt to isolate, measure or control for these variables, and include the Post-Ventilation segment in our analysis simply as a reference period of increased fresh air and sensory disturbance.

### EEG results

To test for the effect of elevated CO_2_ concentration upon participants’ EEG, a 4 (frequency) by 5 (electrode region) by 3 (analysis segment) by 2 (group) mixed model ANOVA was run. Results found a main effect of frequency (*F* (1.08, 36.58) = 89.62, *p* < .001,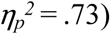, electrode region (*F* (1.50, 51.13) = 50.52, *p* < .001, 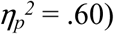, and analysis segment (*F* (2, 68) = 7.98, *p*= .001, 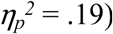In addition significant interactions were also found for frequency by region (*F* (1.72, 58.56) = 34.57, *p* < .001, 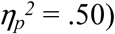frequency by analysis segment (*F* (2.09, 70.95) = 9.16, *p* < .001, 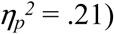region by analysis segment (*F* (2.98, 101.29) = 7.61, *p* < .001, 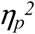 = .18), and frequency by region by analysis segment (*F* (3.73, 126.84) = 4.91, *p* = .001, 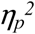 = .13). There was no main effect of group, and no significant group interactions.

Post-hoc analysis of the main effects (Sidak) showed that each frequency significantly differed from the others (*p’s* < .004) such that high-delta had the highest power, followed by theta, then alpha, then beta. Frontal electrodes had greater power than all other regions (*p’s* < .001). Central and temporal electrodes did not differ from each other and neither did parietal and occipital electrodes. Frequency power during Baseline was significantly lower than during the High-CO_2_ (*p* = .001) analysis segment, but did not differ from Post-Ventilation. There was a trend toward the Post-Ventilation analysis segment having a lower overall power than the High-CO_2_ segment (*p* = .09).

To investigate the significant interactions, paired-sample *t*-tests were computed between the Baseline and High-CO_2_ analysis segments and the High-CO_2_ and Post-Ventilation analysis segments for each brain region and frequency, overall and for the TG and CG participants separately (Table 4).

**Table 4:**
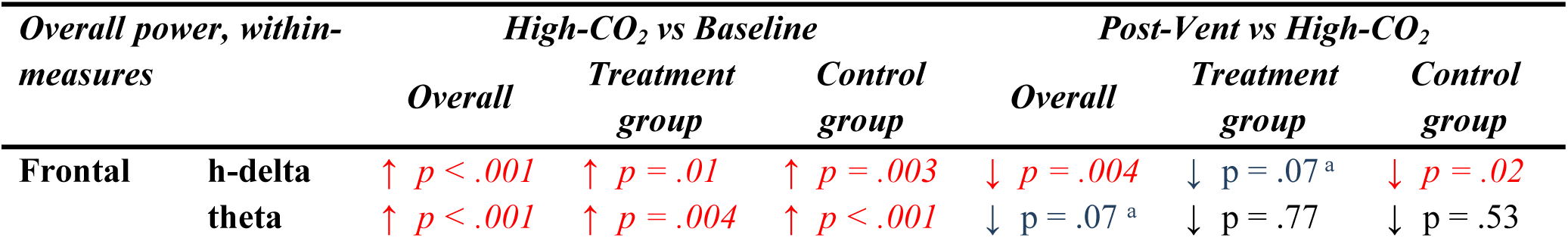
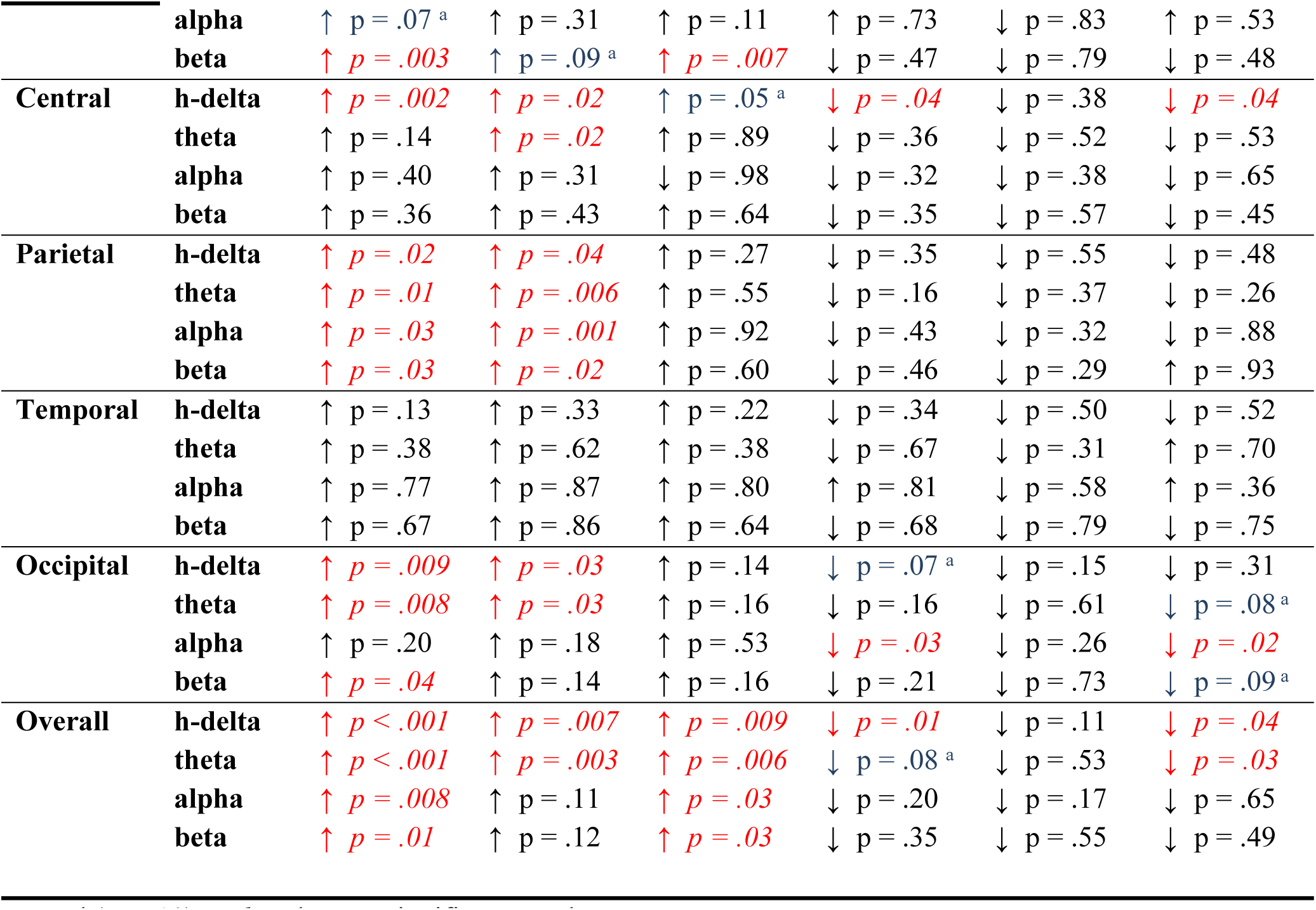
Overall power, within measures, comparison of changes in power by analysis segment for each group. p-values derived from paired sample post-hoc t-tests

| Overall power, within-measures |  | High-CO<sub>2</sub> vs Baseline |  |  | Post-Vent vs High-CO<sub>2</sub> |  |  |
| --- | --- | --- | --- | --- | --- | --- | --- |
|  |  | Overall | Treatment group | Control group | Overall | Treatment group | Control group |
| Frontal | h-delta | ↑ $p < .001$ | ↑ $p = .01$ | ↑ $p = .003$ | ↓ $p = .004$ | ↓ $p = .07^a$ | ↓ $p = .02$ |
| | theta | ↑ $p < .001$ | ↑ $p = .004$ | ↑ $p < .001$ | ↓ $p = .07^a$ | ↓ $p = .77$ | ↓ $p = .53$ |
|  | **alpha** | ↑ *p* = .07<sup>a</sup> | ↑ *p* = .31 | ↑ *p* = .11 | ↑ *p* = .73 | ↓ *p* = .83 | ↑ *p* = .53 |
|  | **beta** | ↑ *p* = .003 | ↑ *p* = .09<sup>a</sup> | ↑ *p* = .007 | ↓ *p* = .47 | ↓ *p* = .79 | ↓ *p* = .48 |
| **Central** | **h-delta** | ↑ *p* = .002 | ↑ *p* = .02 | ↑ *p* = .05<sup>a</sup> | ↓ *p* = .04 | ↓ *p* = .38 | ↓ *p* = .04 |
|  | **theta** | ↑ *p* = .14 | ↑ *p* = .02 | ↑ *p* = .89 | ↓ *p* = .36 | ↓ *p* = .52 | ↓ *p* = .53 |
|  | **alpha** | ↑ *p* = .40 | ↑ *p* = .31 | ↓ *p* = .98 | ↓ *p* = .32 | ↓ *p* = .38 | ↓ *p* = .65 |
|  | **beta** | ↑ *p* = .36 | ↑ *p* = .43 | ↑ *p* = .64 | ↓ *p* = .35 | ↓ *p* = .57 | ↓ *p* = .45 |
| **Parietal** | **h-delta** | ↑ *p* = .02 | ↑ *p* = .04 | ↑ *p* = .27 | ↓ *p* = .35 | ↓ *p* = .55 | ↓ *p* = .48 |
|  | **theta** | ↑ *p* = .01 | ↑ *p* = .006 | ↑ *p* = .55 | ↓ *p* = .16 | ↓ *p* = .37 | ↓ *p* = .26 |
|  | **alpha** | ↑ *p* = .03 | ↑ *p* = .001 | ↑ *p* = .92 | ↓ *p* = .43 | ↓ *p* = .32 | ↓ *p* = .88 |
|  | **beta** | ↑ *p* = .03 | ↑ *p* = .02 | ↑ *p* = .60 | ↓ *p* = .46 | ↓ *p* = .29 | ↑ *p* = .93 |
| **Temporal** | **h-delta** | ↑ *p* = .13 | ↑ *p* = .33 | ↑ *p* = .22 | ↓ *p* = .34 | ↓ *p* = .50 | ↓ *p* = .52 |
|  | **theta** | ↑ *p* = .38 | ↑ *p* = .62 | ↑ *p* = .38 | ↓ *p* = .67 | ↓ *p* = .31 | ↑ *p* = .70 |
|  | **alpha** | ↑ *p* = .77 | ↑ *p* = .87 | ↑ *p* = .80 | ↑ *p* = .81 | ↓ *p* = .58 | ↑ *p* = .36 |
|  | **beta** | ↑ *p* = .67 | ↑ *p* = .86 | ↑ *p* = .64 | ↓ *p* = .68 | ↓ *p* = .79 | ↓ *p* = .75 |
| **Occipital** | **h-delta** | ↑ *p* = .009 | ↑ *p* = .03 | ↑ *p* = .14 | ↓ *p* = .07<sup>a</sup> | ↓ *p* = .15 | ↓ *p* = .31 |
|  | **theta** | ↑ *p* = .008 | ↑ *p* = .03 | ↑ *p* = .16 | ↓ *p* = .16 | ↓ *p* = .61 | ↓ *p* = .08<sup>a</sup> |
|  | **alpha** | ↑ *p* = .20 | ↑ *p* = .18 | ↑ *p* = .53 | ↓ *p* = .03 | ↓ *p* = .26 | ↓ *p* = .02 |
|  | **beta** | ↑ *p* = .04 | ↑ *p* = .14 | ↑ *p* = .16 | ↓ *p* = .21 | ↓ *p* = .73 | ↓ *p* = .09<sup>a</sup> |
| **Overall** | **h-delta** | ↑ *p* < .001 | ↑ *p* = .007 | ↑ *p* = .009 | ↓ *p* = .01 | ↓ *p* = .11 | ↓ *p* = .04 |
|  | **theta** | ↑ *p* < .001 | ↑ *p* = .003 | ↑ *p* = .006 | ↓ *p* = .08<sup>a</sup> | ↓ *p* = .53 | ↓ *p* = .03 |
|  | **alpha** | ↑ *p* = .008 | ↑ *p* = .11 | ↑ *p* = .03 | ↓ *p* = .20 | ↓ *p* = .17 | ↓ *p* = .65 |
|  | **beta** | ↑ *p* = .01 | ↑ *p* = .12 | ↑ *p* = .03 | ↓ *p* = .35 | ↓ *p* = .55 | ↓ *p* = .49 |

<sup>a</sup> Trend ( $p < .10$ ). *Italics* denotes significant *p*-values

Overall results, irrespective of group, show no changes in the temporal electrode region for any frequency. The strongest effects from Baseline to High-CO_2_ are an increase of frontal high-delta, theta and beta, central high-delta, and occipital high-delta and theta, as well as global increases in high-delta, theta, and alpha. Despite a lack of significant group effects in the overall model, the data presented in Table 4 show a clear difference in the pattern of frequency power changes across the brain in the two groups. According to the definition of drowsiness employed (Section 0), the results show the EEG of the TG shows a closer approximation to drowsiness compared to that of the CG, considering: (a) the increase in delta and theta is more global than the CG and (b) CG also has a significant overall increase in alpha and beta, while TG increase is theta and high-delta only.

### Relationship between EEG and temperature

In order to assess whether any relationship existed between the temperature in the room and the EEG, Pearson correlations were run for each analysis segment. The results show no significant correlation between the average temperature during the segment and the global EEG power of each frequency recorded during that time period. Correlations were also run for each electrode region. This analysis found a significant negative relationship for alpha power in the temporal region and temperature during Baseline only (*r* = -.34, *p* = .04).

### Self-reported sleepiness (Effect of analysis segment, treatment group, within measures)

Analysis of questionnaire data on subjective sleepiness found a significant main effect of analysis segment on self-reported sleepiness, *χ*^*2*^ (2) = 22.84, *p* < .001 (Friedman’s ANOVA). Wilcoxon matched pairs post-hoc comparisons show that participants at High-CO_2_ had significantly higher ratings of sleepiness than both Baseline (*p* < .001) and Post-Ventilation (*p*= .01). The Post-Ventilation segment also showed significantly higher ratings of sleepiness than Baseline (*p* = .01) (Table 5). These p-values remained significant when analysed using parametric statistics (3-way ANOVA).

**Table 5:** Self reported sleepiness, average rating with SD, within measures, TG and CG participants

| *Self-reported Sleepiness, Average Rating ± SD, within-measures* |  |  |
| --- | --- | --- |
|  | *Treatment group* | *Control group* |
| **Baseline** | 2.2 ± .7 | 2.2 ± .8 |
| **High-CO<sub>2</sub>** | 3.2 ± 1.1 | 3.7 ± 1.0 |
| **Post-Ventilation** | 2.7 ± 1.2 | 2.6 ± 1.0 |

The average sleepiness ratings are similar for both TG and CG participants; *p* > .05 for both parametric and non-parametric comparisons (Table 5), indicating that subjective sleepiness was not affected by the changes in CO_2_ concentration. None of the group comparisons for sleepiness approach significance.

## Discussion

The effect of office-realistic changes to CO_2_ on resting EEG represent a knowledge gap in the literature to date. This study tests the effect of a 2,700 ppm concentration of CO_2_ in an office on resting EEG, analysing EEG results for indicators of a progression towards drowsiness. Data was analysed at three segments of each experiment; Baseline, High-CO_2_ and Post-Ventilation. This study supports the role of EEG as a means of objectively measuring drowsiness in humans when affected by changes to the indoor climate.

### Evidence for the effect of CO_2_ on drowsiness- Relationship between TG and CG participants’ EEG

Results from this study provide an indication that the indoor CO_2_ concentration of 2,700 ppm had an effect on the EEG indicative of a progression towards drowsiness, when drowsiness is characterised by a global increase in delta and theta [22,23]. Despite the lack of a significant effect of group in the overall model, and both groups showing some evidence of a progression towards drowsiness, the evidence of drowsiness is stronger for the TG (Table 4). A distinct trend observed among TG participants is the global nature of the high-delta and theta increases from Baseline to High-CO_2_ among TG participants relative to the only frontal increase in these parameters among CG participants. The findings of this paper reinforce calls for sufficient ventilation in knowledge work spaces [2] and greater occupant awareness of indoor CO_2_ concentration in these spaces [48].

The Post-Ventilation findings show further differences between the TG and CG, where the CG participants appeared better able to overcome the increased (EEG-assessed) drowsiness experienced in the High-CO_2_ analysis segment. This may imply that the increased CO_2_experienced by TG participants affected the return of the EEG signals to Baseline levels. However given the difference in sample size between the groups, caution must be taken when looking at any potential group differences until further research is conducted with larger, more equal group sizes.

### Relationship between self-reported and EEG-measured drowsiness

The EEG of the TG more closely approximates drowsiness at High CO_2_ compared to the CG. Yet the difference between average self-reported sleepiness ratings at High CO_2_ between CG and TG is minimal (half the standard deviation), and is not significantly different between groups (p > 0.5), (Table 5). Longer exposures to comparable concentrations of CO_2_ with bio- effluents are found to affect (subjectively assessed) drowsiness: 255 minutes exposure to 3,000 ppm with bio effluents increased subjective sleepiness and difficulty in thinking clearly [10]; 235 minutes exposure to 2,260 ppm affected perceived fatigue and perceived lack of energy [4] and four hours’ exposure to CO_2_ above 2,700 ppm resulted in increased subjective sleepiness [14]. The duration of this present study is much shorter than other studies and subjective sleepiness between groups was unaffected. Given the short duration of the study and the similarity of subjective sleepiness between groups, a possible explanation here is that both groups self-report higher feelings of sleepiness simply as a function of time (being sat still in the same room with no stimulation).

Further work is required to determine whether the objectively measured drowsiness indicated in the EEG results persist over longer timescales, whether self-reported drowsiness is better correlated to EEG over time, and whether EEG may be used as something of an early warning system for drowsiness. Small changes in CO_2_ can quickly affect blood pH [31], and owing to the short duration of the experiment, it is possible that EEG results may provide a more timely indication of physiological changes than subjective sleepiness, though this suggestion needs to be corroborated. Additionally, because both subjectively and objectively measured indications of drowsiness were reduced following ventilation of the room future work could additionally explore the potential of regular ventilation episodes in knowledge work spaces to retain alertness.

### Relationship between EEG and temperature

Results also show a significant effect of temperature with CG participants, completing the experiment at a slightly higher temperature than TG participants. Temperature in both groups increased from Baseline to High-CO_2_ before dropping to below baseline levels as a result of the ventilation of the room. Related literature finds lower temperatures (without increased CO_2_) are correlated to decreased drowsiness as measured by EEG [29], and increasing indoor temperatures (i.e. warm discomfort) is correlated to difficulty concentrating [49]. These findings might explain the higher subjective sleepiness experienced by the CG at High CO_2_; however, as mentioned, the subjective sleepiness ratings were small and not statistically significant and all participants were invited to modify their clothing if required in order to remain thermally comfortable throughout the experiment. Conversely, the TG had a higher objective indication of drowsiness but were subject to cooler temperatures than the CG, potentially suggestive that (1) the effects on the EEG of the TG in this study may be attributable to CO_2_ rather than temperature and (2) that subjective and objective determinations of drowsiness may not be correlated over short timescales. Future research could better control the temperature of the environment to remove this variable as a potential confound. Additionally, the correlation between objectively and subjectively measured drowsiness due to changed CO_2_ conditions needs to be further explored, e.g. the potential for EEG to act as an early warning system for drowsiness.

### Limitations and confounding factors

The results of this study should be viewed in light of its limitations: (1) The duration of exposure in this study is much shorter than comparable studies of office-realistic CO_2_ concentrations on humans [8,10,14,16,50], and future work is required to determine whether the changes in EEG with respect to drowsiness are momentary or sustained. (2) Accordingly, changes in the EEG of the TG should be considered as indicative of a neurological progression towards drowsiness, rather than definitive drowsiness. (2) While the CO_2_ outlet was attached to a fan, mixing may not have been as effective as is possible in a climate chamber. (3) All participants assumed that gas was released into the room during the experiment, as the CG participants were exposed to a pre-recorded and equalized sound to mimic the CO_2_ gas being released throughout Condition 3. Thus the participants were blind to the conditions, but were not blinded to the fact that the air in the room was (supposedly) being modified. Thus it cannot be ruled out that some CG may have experienced a placebo reaction. (4) The treatment and control groups differ in sample size and the study is underpowered with respect to between-groups analysis (a-priory power analysis N = 58, i.e. 29 per group), potentially explaining the lack of group effects found in the overall ANOVA. However, even after discarding participants with poor EEG data, the study is still well powered to make conclusions based on the within subjects analysis (a-priori power analysis n = 18) of the whole sample, and for the TG. As such, we are confident in our conclusion that the pattern of results found for this group more closely approximates drowsiness. The study is only slightly under powered with regards to within subjects analysis for the CG group only.

### Future work

To corroborate our findings, future work using EEG as an objective indicator of the effects of changes to indoor air quality would be helpful. To better isolate CO_2_ as a variable in future studies, we suggest a within subjects study design for future work in order to ensure equal representation in the high and “sham” CO_2_ groups. Such a design would control for any individual differences between the groups. Fully blinding participants to experimental conditions might also be beneficial. In addition, there are personal factors not controlled for in this study which could feasibly influence drowsiness, such as number of hours sleep, amount of time since their last meal, their previous activity before experiment. Future studies should account for such factors. Given our finding that a 10 minute ventilation period appeared to reverse the trend towards drowsiness (Post-Vent versus High CO_2_), we suggest further work investigates the acceptability of periodic drafts in naturally ventilated workplaces as a means of maintaining vigilance and concentration.

## Conclusion

Drowsiness represents an important factor affecting office work and productivity [14,20], yet many studies assessing the effects of poor indoor environment quality on humans gather only subjective data for factors potentially affecting work performance such as drowsiness or mood. In this study we have demonstrated the potential for EEG to be used as an objective measurement of drowsiness to determine the effect of elevated levels of indoor CO_2_. Results indicate that even short exposure to elevated levels of CO_2_ indoors (TG) can produce EEG indicative of a progression towards drowsiness. Further work is necessary to corroborate these findings.

Priorities for further work have been outlined including: longer-duration studies using EEG, full blinding to test conditions, accounting for other potential physiological factors which may affect drowsiness (e.g. including time since last meal, hours of sleep), and the acceptability of periodic drafts in naturally ventilated workplaces as a means of maintaining vigilance and concentration.

## Acknowledgements

This work was carried out as part of the Refresh project funded by the Engineering and Physical Sciences Research Council (EPSRC) Grants: EP/K021907/1, EP/K021834/1 and EP/K021893/1. Thanks are required for John Lally for technical assistance. All data supporting this study are openly available from the University of Southampton repository: (DOI TO BE INSERTED ONCE PUBLISHED).

## References

1. Cui S, Cohen M, Stabat P, Marchio D. CO2 tracer gas concentration decay method for measuring air change rate. Build Environ. 2015;84:162–9.

2. Fisk WJ. The ventilation problem in schools: literature review. Vol. 27, Indoor Air. 2017. p. 1039–51.

3. Etheridge D. A perspective on fifty years of natural ventilation research. Build Environ. 2015;91:51–60.

4. Maula H, Hongisto V, Naatula V, Haapakangas A, Koskela H. The effect of low ventilation rate with elevated bioeffluent concentration on work performance, perceived indoor air quality, and health symptoms. Indoor Air. 2017;27(6):1141–53.

5. Mendell MJ, Heath GA. Do indoor pollutants and thermal conditions in schools influence student performance? A critical review of the literature. Indoor Air. 2005;15(1):27–52.

6. Wargocki P, Sundell J, Bischof W, Brundrett G, Fanger PO, Gyntelberg F, et al. Ventilation and health in non-industrial indoor environments: Report from a European Multidisciplinary Scientific Consensus Meeting (EUROVEN). Vol. 12, Indoor Air. 2002. p. 113–28.

7. Wyon DP. The effects of indoor air quality on performance and productivity. Indoor Air. 2004;14 Suppl 7(Suppl 7):92–101.

8. Maddalena R, Mendell MJ, Eliseeva K, Chan WR, Sullivan DP, Russell M, et al. Effects of ventilation rate per person and per floor area on perceived air quality, sick building syndrome symptoms, and decision-making. Indoor Air. 2015;25(4):362–70.

9. Fang L, Wyon DP, Clausen G, Fanger PO. Impact of indoor air temperature and humidity in an office on perceived air quality, SBS symptoms and performance. Indoor Air, Suppl. 2004;14(SUPPL. 7):74–81.

10. Zhang X, Wargocki P, Lian Z, Thyregod C. Effects of exposure to carbon dioxide and bioeffluents on perceived air quality, self-assessed acute health symptoms, and cognitive performance. Indoor Air. 2017;27(1):47–64.

11. Wargocki P, Wyon DP. Ten questions concerning thermal and indoor air quality effects on the performance of office work and schoolwork. Build Environ. 2017;112:359–66.

12. Zhang X, Wargocki P, Lian Z. Human responses to carbon dioxide, a follow-up study at recommended exposure limits in non-industrial environments. Build Environ. 2016;100:162–71.

13. Zhang X, Wargocki P, Lian Z. Effects of Exposure to Carbon Dioxide and Human Bioeffluents on Cognitive Performance. Procedia Eng. 2015;121:138–42.

14. Vehviläinen T, Lindholm H, Rintamäki H, Pääkkönen R, Hirvonen A, Niemi O, et al. High indoor CO 2 concentrations in an office environment increases the transcutaneous CO 2 level and sleepiness during cognitive work. J Occup Environ Hyg. 2016;13(1):19–29.

15. Allen JG, MacNaughton P, Satish U, Santanam S, Vallarino J, Spengler JD. Associations of cognitive function scores with carbon dioxide, ventilation, and volatile organic compound exposures in office workers: A controlled exposure study of green and conventional office environments. Environ Health Perspect. 2016;124(6):805–12.

16. Satish U, Mendell MJ, Shekhar K, Hotchi T, Sullivan D, Streufert S, et al. Is CO2 an indoor pollutant? Direct effects of low-to-moderate CO2 concentrations on human decision-making performance. Environ Health Perspect. 2012 Sep 20;120(12):1671–7.

17. Kajtár L, Herczeg L. Influence of carbon-dioxide concentration on human well-being and intensity of mental work. Idojaras. 2012;116(2):145–69.

18. Liu W, Zhong W, Wargocki P. Performance, acute health symptoms and physiological responses during exposure to high air temperature and carbon dioxide concentration. Build Environ. 2017;114:96–105.

19. Zhang X, Wargocki P, Lian Z. Physiological responses during exposure to carbon dioxide and bioeffluents at levels typically occurring indoors. Indoor Air. 2017;27(1):65–77.

20. Tanabe S, Nishihara N. Productivity and fatigue. Indoor Air, Suppl. 2004 Aug 1;14(SUPPL. 7):126–33.

21. Lal skl, Craig A. Driver fatigue: Electroencephalography and psychological assessment. Psychophysiology. 2002;39(3):313–21.

22. De Gennaro L, Marzano C, Veniero D, Moroni F, Fratello F, Curcio G, et al. Neurophysiological correlates of sleepiness: A combined TMS and EEG study. Neuroimage. 2007;39(3):313–21.

23. Gorgoni M, Ferlazzo F, Ferrara M, Moroni F, D’Atri A, Fanelli S, et al. Topographic electroencephalogram changes associated with psychomotor vigilance task performance after sleep deprivation. Sleep Med. 2014;15(9):1132–9.

24. Trejo LJ, Kubitz K, Rosipal R, Kochavi RL, Montgomery LD, Rosepal R, et al. EEG- Based Estimation and Classification of Mental Fatigue. Psychology. 2015;(April):1–44.

25. Wargocki P, Wyon DP, Sundell J, Clausen G, Fanger PO. The Effects of Outdoor Air Supply Rate in an Office on Perceived Air Quality, Sick Building Syndrome (SBS) Symptoms and Productivity. Indoor Air. 2000;10(4):222–36.

26. Donaldson SI, Grant-Vallone EJ. Understanding self-report bias in organizational behavior research. J Bus Psychol. 2002;17(2):245–60.

27. Rasheed EO, Byrd H. Can self-evaluation measure the effect of IEQ on productivity? A review of literature. Facilities. 2017;35(11/12):601–21.

28. Niedermeyer E, Silva FHL Da. Electroencephalography: Basic Principles, Clinical Applications, and Related Fields. 1st ed. Lippincott Williams and Wilkins. Lippinciott Williams and Wilkins; 2004. 1309 p.

29. Landstrom U, Englund K, Nordstrom B, Stenudd A. Laboratory studies on the effects of temperature variations on drowsiness. Percept Mot Skills. 1999;83(3):1217–29.

30. Wang D, Yee BJ, Wong KK, Kim JW, Dijk DJ, Duffin J, et al. Comparing the effect of hypercapnia and hypoxia on the electroencephalogram during wakefulness. Clin Neurophysiol. 2015;126(1):103–9.

31. Xu F, Uh J, Brier MR, Hart J, Yezhuvath US, Gu H, et al. The influence of carbon dioxide on brain activity and metabolism in conscious humans. J Cereb Blood Flow Metab. 2011;31(1):58–67.

32. Thesen T, Leontiev O, Song T, Dehghani N, Hagler DJ, Huang M, et al. Depression of cortical activity in humans by mild hypercapnia. Hum Brain Mapp. 2012;33(3):715–26.

33. Bloch-Salisbury E, Lansing R, Shea S a. Acute changes in carbon dioxide levels alter the electroencephalogram without affecting cognitive function. Psychophysiology. 2000;37(4):418–26.

34. Iber C, Ancoli-Israel S, Chesson AL Jr. QS for the aa, Medicine. of S, Iber CC, Ancoli-Israel S, Chesson AL, Quan SF, et al. The AASM Manual for the Scoring of Sleep and Associated Events: Rules, Terminology and Technical Specifications. In: Berry, R.B., Brooks, R., Gamaldo, C.E., Harding, S.M., Marcus, C.L. and Vaughn BV, editor. AASM Manual for Scoring Sleep. Westchester, IL; 2007.

35. Lamb S, Kwok kcs. A longitudinal investigation of work environment stressors on the performance and wellbeing of office workers. Appl Ergon. 2016;52:104–11.

36. Garner M, Attwood A, Baldwin DS, Munafò MR. Inhalation of 7.5% carbon dioxide increases alerting and orienting attention network function. Psychopharmacology (Berl). 2012;223(1):67–73.

37. Faul F, Erdfelder E, Lang A-G, Buchner A. G*Power 3: A flexible statistical power analysis program for the social, behavioral, and biomedical sciences. Behav Res Methods. 2007;39(2):175–91.

38. Laussmann D, Helm D. Air Change Measurements Using Tracer Gases. Chem Emiss Control Radioact Pollut Indoor Air Qual. 2004;1:365–404.

39. Gough HL, Luo Z, Halios CH, King MF, Noakes CJ, Grimmond csb, et al. Field measurement of natural ventilation rate in an idealised full-scale building located in a staggered urban array: Comparison between tracer gas and pressure-based methods. Build Environ. 2018;137:246–56.

40. Hedge A, Erickson WA. A study of indoor environment and sick building syndrome complaints in air-conditioned offices: benchmarks for facility performance. Int J Facil Manag. 1997;1(4):185–92.

41. Watson D, Clark LA, Tellegen A. Development and validation of brief measures of positive and negative affect: The PANAS scales. J Pers Soc Psychol. 1988;54(6):1063–70.

42. Quest S. Stanford Sleepiness Scale [Internet]. Stanford Sleepiness Scale. 2000 [cited 2017 Jan 31]. p. 1. Available from: https://web.stanford.edu/∼dement/sss.html

43. American Society of Heating Refrigerating and Air Conditioning Engineers (ASHRAE). ANSI/ASHRAE Standard 55: Thermal Environmental Conditions for Human Occupancy. [Internet]. ASHRAE. 2013. Available from: https://www.ashrae.org/technical-resources/bookstore/thermal-environmental-conditions-for-human-occupancy

44. Biswas D, Bono V, Scott-South M, Chatterjee S, Soska A, Snow S, et al. Analysing wireless EEG based functional connectivity measures with respect to change in environmental factors. In: 3rd IEEE EMBS International Conference on Biomedical and Health Informatics, BHI 2016. 2016. p. 599–602.

45. Bono V, Jamal W, Das S, Maharatna K. Artifact reduction in multichannel pervasive EEG using hybrid WPT-ICA and WPT-EMD signal decomposition techniques. In: ICASSP, IEEE International Conference on Acoustics, Speech and Signal Processing - Proceedings. 2014. p. 5864–8.

46. Tatum W, Husain A, Benbadis S, Kaplan P. Handbook of EEG Interpretation. Medicine. New York: Demos Medical Publishing; 2008.

47. Tinguely G, Finelli LA, Landolt HP, Borbély AA, Achermann P. Functional eeg topography in sleep and waking: State-dependent and state-independent features. Neuroimage. 2006;32(1):283–92.

48. Wargocki P, Da Silva naf. Use of visual CO2 feedback as a retrofit solution for improving classroom air quality. Indoor Air. 2015 Feb 1;25(1):105–14.

49. Mendell MJ, Mirer AG. Indoor thermal factors and symptoms in office workers: findings from the us epa base study. Indoor Air. 2009;19(4):291–302.

50. Maula H, Hongisto V, Koskela H, Haapakangas A. The effect of cooling jet on work performance and comfort in warm office environment. Vol. 104, Building and Environment. 2016. p. 13–20.

